# Regulation of perineuronal nets in the adult cortex by the electrical activity of parvalbumin interneurons

**DOI:** 10.1101/671719

**Authors:** Gabrielle Devienne, Sandrine Picaud, Ivan Cohen, Juliette Piquet, Ludovic Tricoire, Damien Testa, Ariel A. Di Nardo, Jean Rossier, Bruno Cauli, Bertrand Lambolez

## Abstract

Perineuronal net (PNN) accumulation around parvalbumin-expressing (PV) inhibitory interneurons marks the closure of critical periods of high plasticity, whereas PNN removal reinstates juvenile plasticity in the adult cortex. Using targeted chemogenetic *in vivo* approaches in the adult mouse visual cortex, we found that transient electrical silencing of PV interneurons, directly or through inhibition of local excitatory neurons, induced PNN regression. Conversely, excitation of either neuron types did not reduce the PNN. We also observed that chemogenetically inhibited PV interneurons exhibited reduced PNN compared to their untransduced neighbors, and confirmed that single PV interneurons express multiple genes enabling cell-autonomous control of their own PNN density. Our results indicate that PNNs are dynamically regulated in the adult by PV neurons acting as sensors of their local microcircuit activities. PNN regulation provides individual PV neurons with an activity-dependent mechanism to control the local remodeling of adult cortical circuits.

## INTRODUCTION

During the post-natal development of the cerebral cortex, the closure of a highly plastic period, called critical period, is concomitant with the accumulation of the PNN, a specialized extracellular matrix enwrapping fast-spiking PV interneurons (Hensch, 2005). The PNN is made of lecticans, proteoglycan link proteins, and tenascin R, it is reticulated and attached to the membrane via hyaluronan, and it can be degraded by various proteases (Dityatev et al., 2010; Kwok et al., 2012; Ferrer-Ferrer and Dityatev, 2018). The PNN attracts in part the homeoprotein transcription factor OTX2 from cerebrospinal fluid to accumulate within PV cells, which in turn enhances PNN accumulation (Sugiyama et al., 2008). Enzymatic digestion of the PNN, modulating the inhibitory tone, or antagonizing OTX2 import by PV cells, reinstate high circuit plasticity in the adult; and a decrease of the PNN accompanies the reopening of plasticity, whatever the paradigm used (Hensch et al., 1998; Pizzorusso et al., 2002; Fagiolini et al., 2004; Beurdeley et al., 2012; Lensjø et al., 2017a; Harauzov et al., 2010; Sale et al., 2010). Conversely, PNN stability is linked to memory resilience, and PNN deficits are thought to contribute to circuit dysfunctions in several pathologies of the central nervous system (Testa et al., 2019).

Alteration of GABAergic transmission can induce PNN regression and reinstate high cortical plasticity, indicating that the PNN is dynamically regulated in the adult (Hensch, 2005; Harauzov et al., 2010; Sale et al., 2010). PV interneurons are strongly interconnected with excitatory pyramidal neurons, express multiple genes involved in PNN synthesis and degradation, and their maturation parallels that of their PNN (Angulo et al., 1999; Ascoli et al., 2008; Okaty et al., 2009; Rossier et al., 2015). This suggests that PV cells are key actors in the physiological regulation of the PNN. Likewise, transient and targeted inhibition of PV cells using chemogenetics (Alexander et al., 2009) *in vivo* is sufficient to restore visual plasticity in the mouse cortex after closure of the critical period (Kuhlman et al., 2013). We hypothesize that this chemogenetic paradigm induces PNN reduction, making the network permissive to circuit plasticity.

Here, we used targeted chemogenetic *in vivo* approaches to test this hypothesis and examine the physiological factors that govern PNN remodeling in the adult mouse visual cortex. We also assessed the acute electrophysiological effects of chemogenetic paradigms. We found that silencing of PV interneurons, directly or through inhibition of excitatory neurons, induced PNN regression, and obtained evidence for cell-autonomous regulation of its own PNN by each PV cell.

## RESULTS

### Targeted chemogenetic inhibition of PV interneurons induces PNN regression in the adult visual cortex

The PNN accumulates postnatally to reach adult density at P50 in the V1 area of the mouse visual cortex (Ye et al., 2018; Lee et al., 2017; Lensjø et al., 2017b). In order to test the hypothesis that targeted chemogenetic inhibition of PV interneurons alters adult PNN density, we adapted the protocol known to reinstate visual plasticity at P35 (Kuhlman et al., 2013) to measure PNN changes between P58 and P61 (see Methods and Fig.S1). Hemilateral injection of Cre-dependent AAV encoding the inhibitory DREADD hM4Di fused to fluorescent protein mCherry in the V1 area of the visual cortex of PV-Cre mice resulted in robust and selective expression in PV interneurons, with 100 % of mCherry+ cells being also PV+ (n=208 out of 208 cells, Fig.S2). Four weeks after viral injection, mice were treated with the DREADD agonist CNO or with PBS. PNNs are most abundant at the peak of PV interneuron distribution in layer IV and upper layer V (Ye et al., 2013; Lensjø et al., 2017b; Rudy et al., 2011). We quantified PNN density around PV+ cells in these layers using WFA staining (Methods and Fig.S1). Following CNO treatment, the PNN of hM4Di-mCherry+ cells was strongly decreased as compared to the PNN of the uninjected contralateral hemicortex (39.7 ± 4.4 % of contralateral PNN density, n=70 mCherry^+^/PV+ cells and n=72 contralateral PV+ cells from 3 animals, p<0.05, Fig.1, Table 1). Conversely, no significant change of the PNN of hM4Di-mCherry+ cells was observed as compared to contralateral PNN in PBS treated animals (101.1 ± 8.5 % of contralateral PNN density, n=85 mCherry^+^ cells and n=87 contralateral cells from 3 animals, p=0.3, Fig.1, Table 1). The significantly lower PNN density of hM4Di+ cells from CNO-treated animals compared to hM4Di+ cells from PBS-treated animals (Table 1) is illustrated in Fig.1. These results indicate that activation by CNO of hM4Di expressed in PV interneurons locally induces PNN regression in the adult visual cortex.

**Figure 1.**
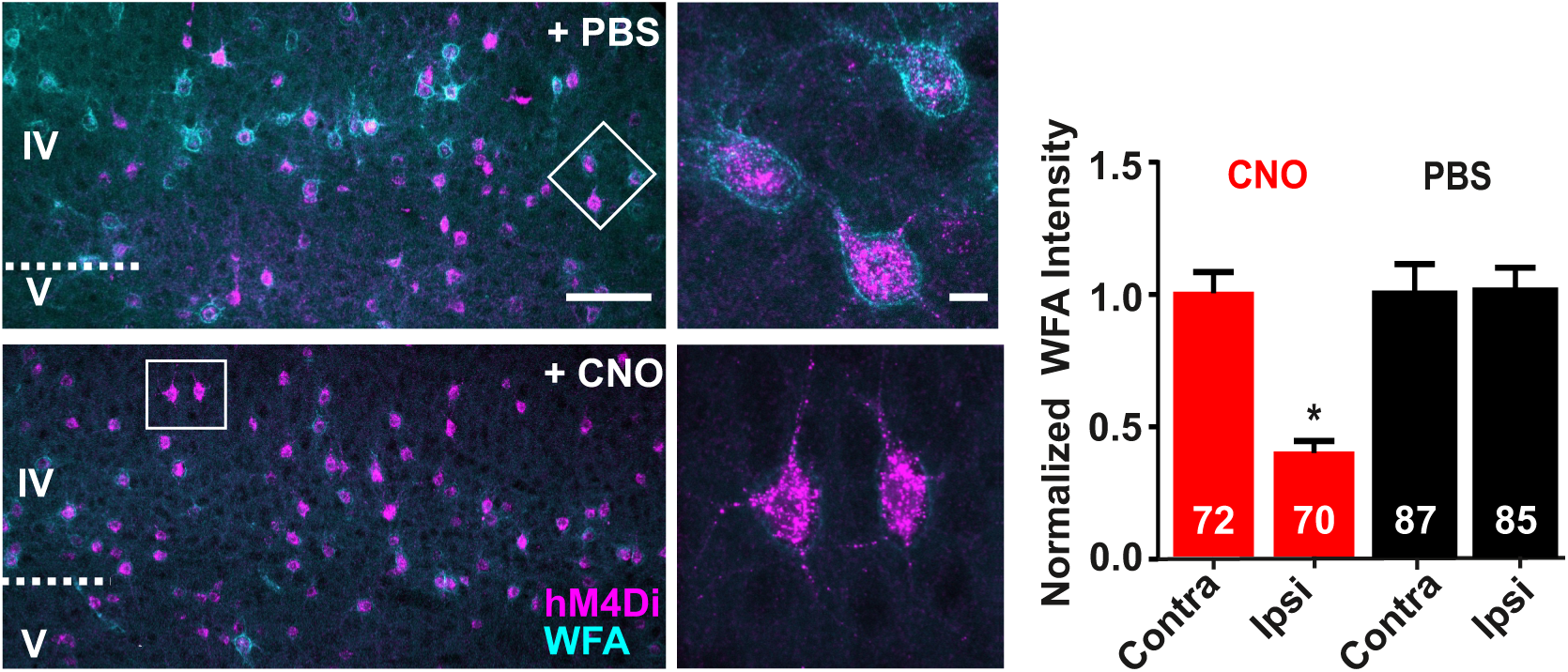
Targeted inhibition of PV interneurons using DREADD hM4Di induces PNN regression. Confocal fluorescence images acquired in the V1 cortex illustrate the PNN (WFA) surrounding hM4Di-expressing PV interneurons in layers IV-V after PBS or CNO treatment of the mice. Note the low PNN density around hM4Di+ cells after CNO treatment, as exemplified in high magnification images. Scale bars: 100 µm (left), 10 µm (right). Plot of PNN density around PV+ (contralateral uninjected hemicortex) and hM4Di+/PV+ (ipsilateral injected) cells in the V1 cortex normalized for each mouse to mean density in contralateral hemicortex. Indicated in bars are the number of cells analyzed in 3 CNO-treated and 3 PBS-treated mice. * Significantly different from other conditions.

**Table 1.**
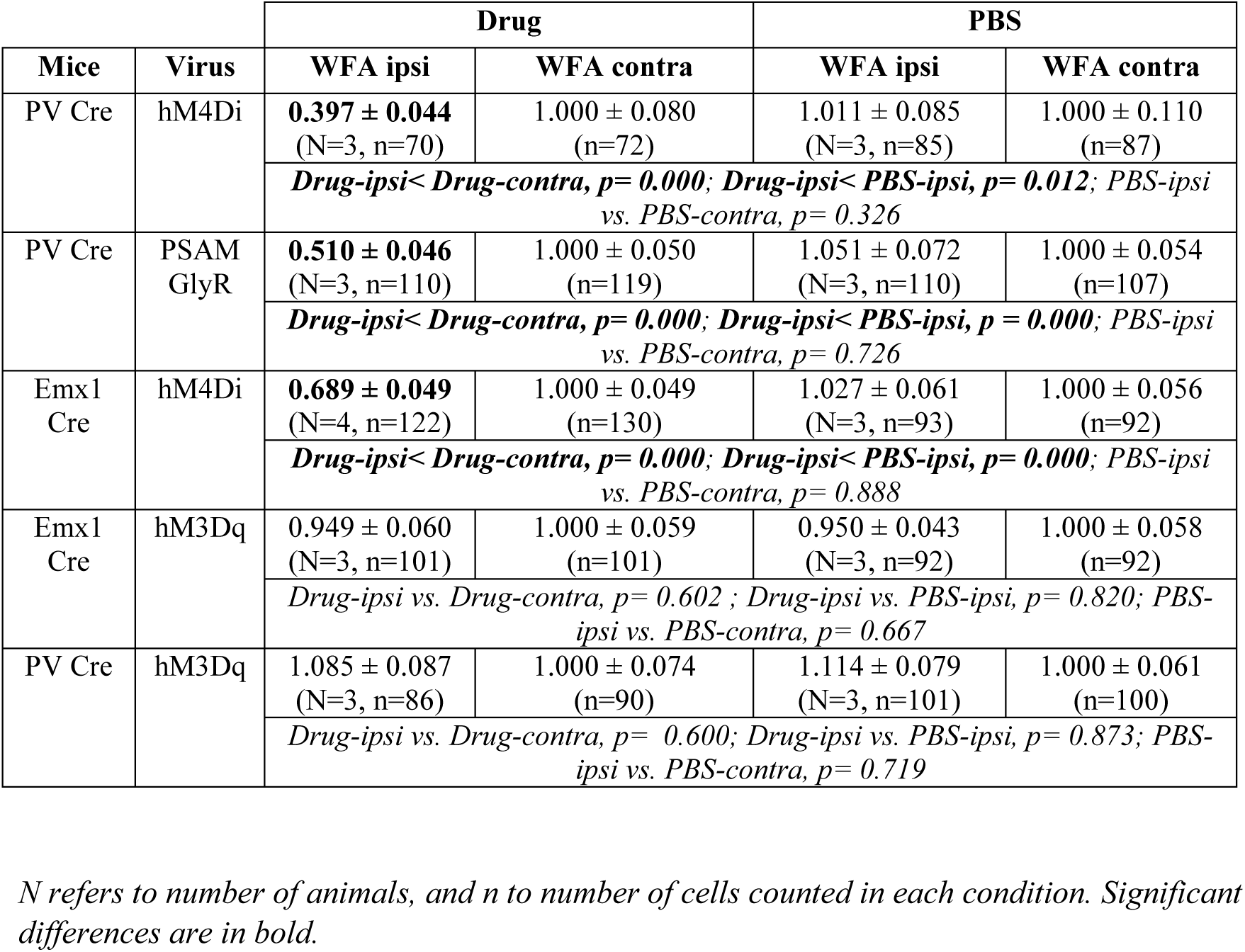
Summary of chemogenetic experiments

| Mice | Virus | Drug |  | PBS |  |
| --- | --- | --- | --- | --- | --- |
|  |  | WFA ipsi | WFA contra | WFA ipsi | WFA contra |
| PV Cre | hM4Di | **0.397 ± 0.044**<br>(N=3, n=70) | 1.000 ± 0.080<br>(n=72) | 1.011 ± 0.085<br>(N=3, n=85) | 1.000 ± 0.110<br>(n=87) |
|  |  | *Drug-ipsi < Drug-contra, p = 0.000; Drug-ipsi < PBS-ipsi, p = 0.012; PBS-ipsi vs. PBS-contra, p = 0.326* |  |  |  |
| PV Cre | PSAM GlyR | **0.510 ± 0.046**<br>(N=3, n=110) | 1.000 ± 0.050<br>(n=119) | 1.051 ± 0.072<br>(N=3, n=110) | 1.000 ± 0.054<br>(n=107) |
|  |  | *Drug-ipsi < Drug-contra, p = 0.000; Drug-ipsi < PBS-ipsi, p = 0.000; PBS-ipsi vs. PBS-contra, p = 0.726* |  |  |  |
| Emx1 Cre | hM4Di | **0.689 ± 0.049**<br>(N=4, n=122) | 1.000 ± 0.049<br>(n=130) | 1.027 ± 0.061<br>(N=3, n=93) | 1.000 ± 0.056<br>(n=92) |
|  |  | *Drug-ipsi < Drug-contra, p = 0.000; Drug-ipsi < PBS-ipsi, p = 0.000; PBS-ipsi vs. PBS-contra, p = 0.888* |  |  |  |
| Emx1 Cre | hM3Dq | 0.949 ± 0.060<br>(N=3, n=101) | 1.000 ± 0.059<br>(n=101) | 0.950 ± 0.043<br>(N=3, n=92) | 1.000 ± 0.058<br>(n=92) |
|  |  | *Drug-ipsi vs. Drug-contra, p = 0.602 ; Drug-ipsi vs. PBS-ipsi, p = 0.820; PBS-ipsi vs. PBS-contra, p = 0.667* |  |  |  |
| PV Cre | hM3Dq | 1.085 ± 0.087<br>(N=3, n=86) | 1.000 ± 0.074<br>(n=90) | 1.114 ± 0.079<br>(N=3, n=101) | 1.000 ± 0.061<br>(n=100) |
|  |  | *Drug-ipsi vs. Drug-contra, p = 0.600; Drug-ipsi vs. PBS-ipsi, p = 0.873; PBS-ipsi vs. PBS-contra, p = 0.719* |  |  |  |

*N* refers to number of animals, and *n* to number of cells counted in each condition. Significant differences are in bold.

The DREADD hM4Di has somatodendritic effects, but can also inhibit GABA release from axon terminals of PV interneurons (Alexander et al., 2009; Kruglikov and Rudy, 2008; Stachniak et al., 2014). Hence, PNN regression may result either from decreased somatodendritic excitation of PV interneurons, or from increased network activity due to disinhibition of their excitatory target neurons. To discriminate between these possibilities, we characterized acute electrophysiological effects of CNO in hM4Di-expressing mice.

### CNO treatment results in moderate silencing of hM4Di-expressing PV interneurons

We investigated the effect of CNO [0.5 µM, (Alexander et al., 2009)] on the excitability of hM4Di-expressing PV interneurons using patch-clamp recordings in acute slices of visual cortex (see Methods). Expression of hM4Di did not conspicuously alter electrophysiological properties of PV interneurons in the absence of CNO (Cauli et al., 1997). CNO application elicited a hyperpolarization (from −66.8 ± 0.5 mV in control to −71.6 ± 0.5 mV in CNO, p<0.05) and a decrease in input resistance (from 152.9 ± 13.6 MΩ in control to 128.6 ± 10.7 MΩ in CNO, p<0.05, n=8 PV cells, Fig.2A), as reported (Alexander et al., 2009). This effect was associated with an increase in rheobase (from 143 ± 16 pA in control to 191 ± 23 pA in CNO, p<0.05, Fig.2A). These results indicate that activation of hM4Di by CNO decreases the somatodendritic excitability of PV neurons.

**Figure 2.**
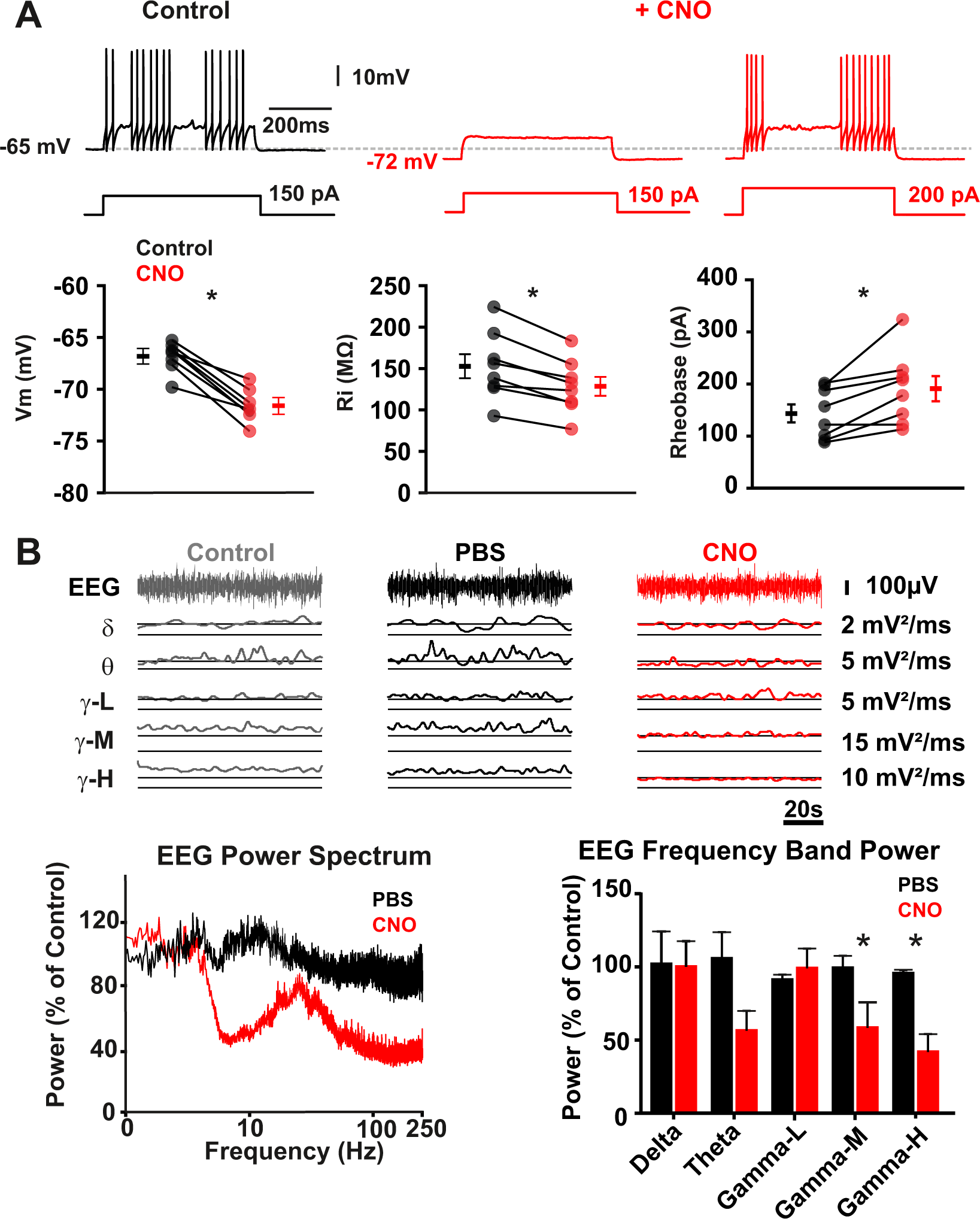
CNO decreases the excitability of hM4Di-expressing PV interneurons and reduces cortical gamma oscillations. (**A**) Patch-clamp recordings in cortical slices. Traces show responses of a hM4Di-expressing interneuron to depolarizing current step in control conditions and upon bath application of CNO (0.5 µM). CNO elicited a hyperpolarization of the membrane potential and an increase in the current needed to induce action potential firing. Plots show parameters measured in hM4Di-expressing interneurons (n=8). Note the large amplitude of the fast afterhyperpolarizing potentials, the quiescent periods between trains of action potentials, the modest input resistance, and the high rheobase value typical of fast-spiking PV interneurons. (**B**) EEG recordings in the hM4Di-expressing visual hemicortex of awake PV-Cre mouse before and after consecutive i.p. injections of PBS and CNO. The EEG signal was filtered to analyze oscillations in the delta 1-4 Hz, theta 6-10 Hz, gamma low 30-50 Hz, gamma mid 55-95 Hz and gamma high 100-150 Hz frequency ranges. Traces show samples obtained from 1 mouse and graphs show mean results from 3 mice. * Significant differences.

We next investigated acute CNO effects on network activities in V1 using EEG recordings in the hemicortex of awake mice expressing hM4Di in PV interneurons (see Methods). After 1 h baseline recording in control conditions, a first i.p. injection of PBS was performed, followed 1 h later by i.p. injection of CNO. We found no evidence for CNO-induced unbalanced excitation of the network indicative of marked disinhibition (Fig.2B). CNO injection induced a strong decrease of network oscillations in the mid-high gamma frequency band (from 99.5 ± 8.5 % of control in PBS to 58.7 ± 17.5 % of control in CNO for the 55-95 Hz range, and from 94.9 ± 2.2 % of control in PBS to 41.1 ± 12.3 % of control in CNO for the 100-150 Hz range, n=3 mice, p<0.05, Fig.2B). We also observed a non-significant decrease in theta oscillations (from 105.7 ± 18.0 % of control in PBS to 56.3 ± 13.6 % of control in CNO, p=0.14, Fig.2B), but no change in low frequency gamma or delta oscillations. PV interneurons play a key role in synchronizing cortical neuronal populations, notably at gamma frequencies (Cardin et al., 2009). Hence, these results indicate that hM4Di activation by CNO induces moderate silencing of PV cells *in vivo*, which could be the trigger of the observed PNN reduction.

### Targeted excitation of glutamatergic neurons or PV interneurons does not alter adult PNN density

In order to rule out disinhibition-induced network excitation as a cause of PNN regression, we selectively expressed the excitatory DREADD hM3Dq in either glutamatergic neurons or PV interneurons using the same paradigm as above in Emx1-Cre or PV-Cre mice, respectively (see Methods). We first verified that CNO (0.5 µM) enhanced the excitability of hM3Dq-expressing neurons using patch-clamp recordings of hM3Dq+ layer V pyramidal cells (n=8) in cortical slices. CNO elicited a depolarization (from −66.1 ± 0.9 mV in control to −63.9 ± 1.4 mV in CNO, p<0.05) and an increase in input resistance (from 139.8 ± 12.5 MΩ in control to 173.2 ± 10.6 MΩ in CNO, p<0.05; Fig.S3A) of these neurons, accompanied by a decrease in rheobase (from 60 ± 3 pA in control to 38 ± 3 pA in CNO, p<0.05; Fig.S3A). These results confirm that CNO enhances the excitability of hM3Dq-expressing neurons.

We next probed the effect of *in vivo* activation of hM3Dq-expressing glutamatergic neurons in V1 on the PNN surrounding PV interneurons. Neither CNO, nor PBS treatment significantly changed PNN density around PV+ cells in the ipsilateral hM3Dq-expressing hemicortex as compared to the uninjected contralateral hemicortex (Fig.S3B, Table 1). We also tested the effect of *in vivo* activation of hM3Dq-expressing PV interneurons in V1 on their PNN. Neither CNO, nor PBS treatment significantly changed the density of the PNN around hM3Dq+/PV+ cells as compared to contralateral PV+ cells (Fig.S3C, Table 1). These results show that cortical network excitation does not trigger PNN regression in the adult. Hence, CNO-induced PNN regression around hM4Di+ PV interneurons did not result from cortical network disinhibition.

### Electrical silencing of PV interneurons triggers PNN regression in the adult visual cortex

The DREADD hM4Di is coupled to Gi intracellular signaling and thus results in both electrophysiological and metabotropic effects. In order to assess electrical silencing of PV interneurons as a cause of PNN decrease, and rule out involvement of Gi-dependent metabotropic effects, we used an alternative chemogenetic tool, the chloride channel PSAM-GlyR exclusively activated by the agonist PSEM^89S^ (Magnus et al., 2011) co-expressed with GFP, to inhibit PV interneurons in the same paradigm as above (see Methods). We found that 82 % of GFP+ cells were PV+ (n=75 out of 92, not shown), consistent with efficient expression of PSAM-GlyR in PV interneurons. Following PSEM^89S^ treatment, the density of the PNN surrounding PSAM-GlyR-GFP+/PV+ cells in layers IV-V was largely decreased as compared to the PNN of contralateral PV+ cells (51.0 ± 4.6 % of contralateral PNN density, n=110 GFP^+^ cells and n=119 contralateral cells from 3 animals, p<0.05; Fig.3, Table 1). PNN density around GFP+ cells did not significantly differ from contralateral PNN in PBS treated animals (Fig.3, Table 1). These results confirm that electrical silencing of PV interneurons induces PNN regression in the adult visual cortex.

**Figure 3.**
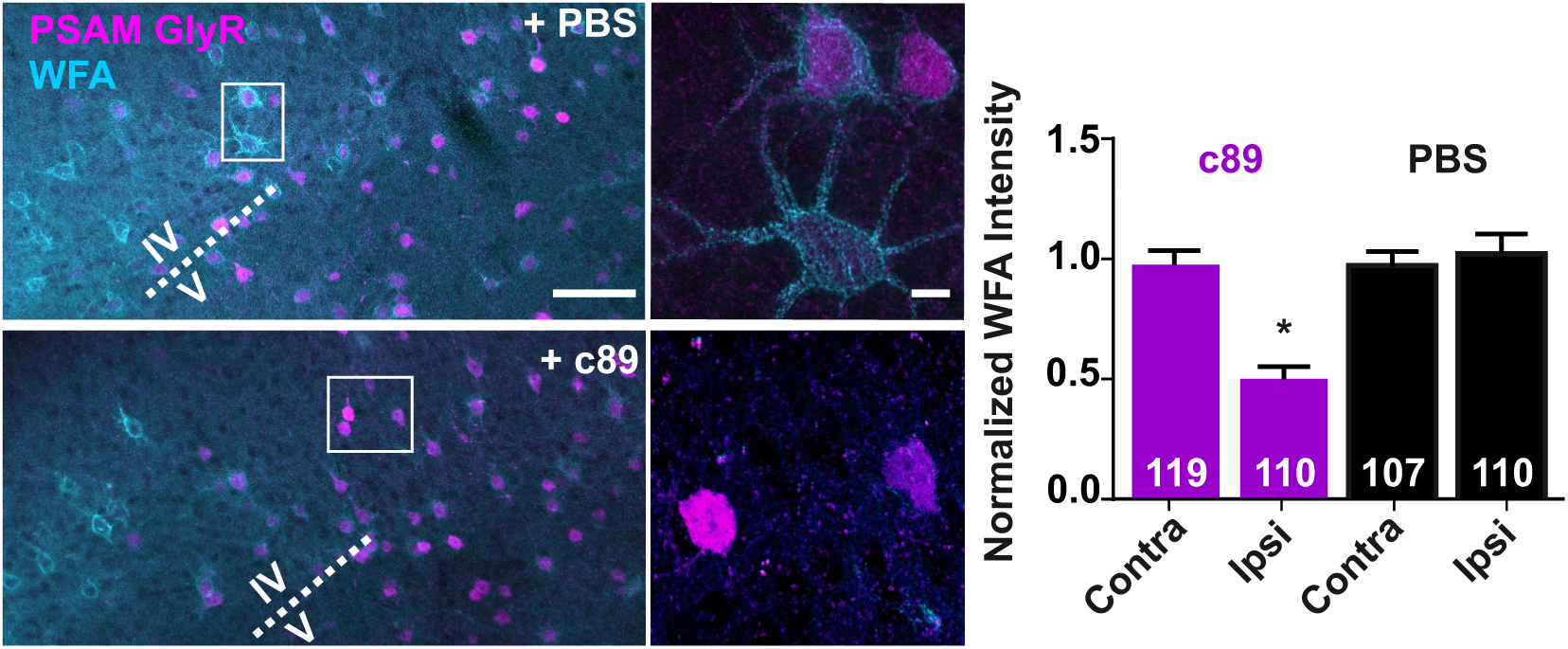
Silencing of PV interneurons by PSAM-GlyR induces PNN regression. Confocal fluorescence images in the V1 cortex illustrate the PNN (WFA) surrounding PSAM-GlyR+ cells after mice treatment with PBS or with the PSAM-GlyR agonist PSEM^89S^. Note the low PNN density around PSAM-GlyR+ cells after PSEM treatment, as exemplified in high magnification images. Scale bars: 100 µm (left), 10 µm (right). Plot of PNN density around PV+ (contralateral uninjected hemicortex) and PSAM-GlyR+/PV+ (ipsilateral injected) cells in the V1 cortex. Indicated in bars are the number of cells analyzed in 3 PSEM-treated and 3 PBS-treated mice. * Significantly different from other conditions.

We next reasoned that PV interneuron silencing can, in principle, also be achieved by decreasing their synaptic excitation. We thus targeted expression of hM4Di to inhibit excitatory neurons in the V1 area. CNO treatment significantly reduced the PNN of PV+ cells in the ipsilateral hM4Di-expressing hemicortex as compared to contralateral PNN (68.9 ± 4.9 % of contralateral PNN density, n=122 ipsilateral and n=130 contralateral PV+ cells from 4 animals, p<0.05; Fig.4, Table 1). PBS treatment had no significant effect on PNN density (Fig.4, Table 1). These data indicate that a decrease in the synaptic excitation of PV interneurons induces PNN regression in the adult visual cortex.

**Figure 4.**
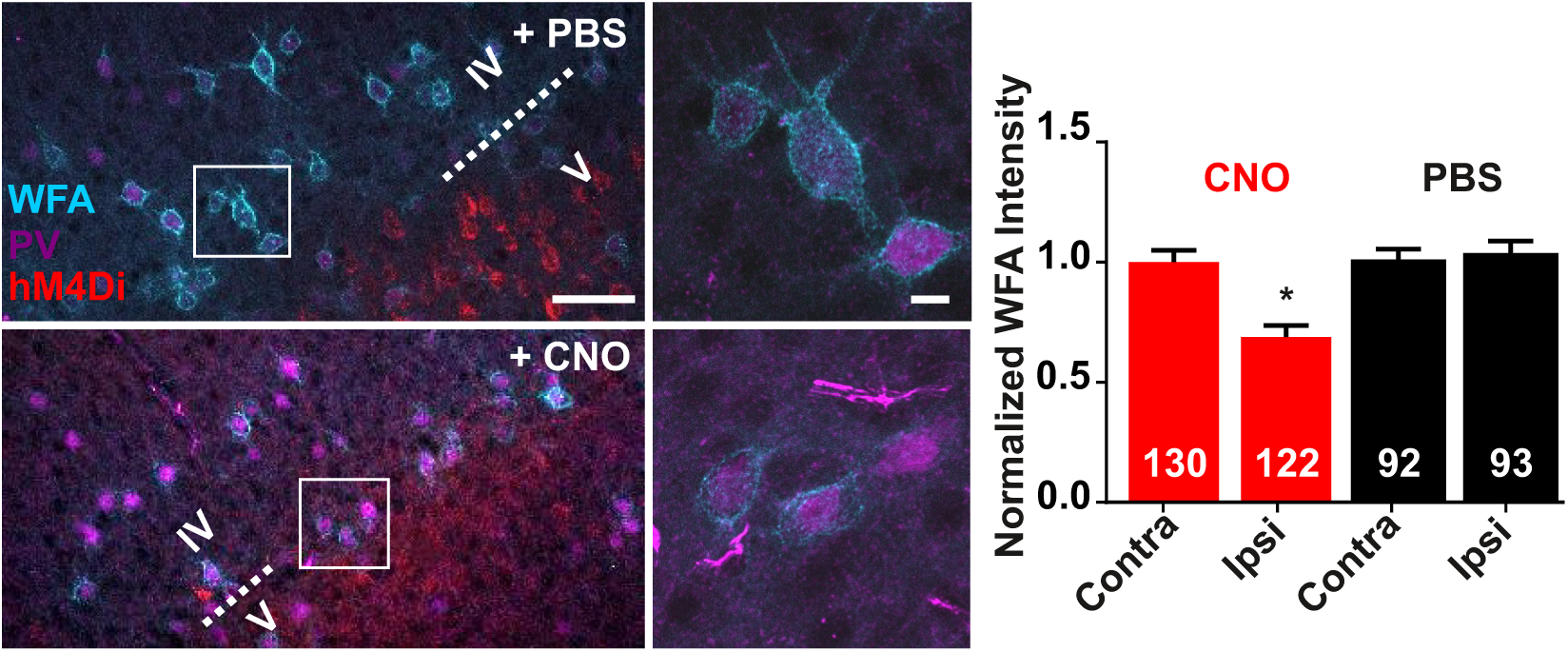
Targeted inhibition of excitatory neurons by hM4Di induces PNN regression. Confocal fluorescence images illustrate the PNN (WFA) surrounding PV+ cells in the vicinity of hM4Di-expressing excitatory neurons in layers IV-V of the V1 cortex of mice treated with PBS or CNO. Note the low PNN density around PV+ cells after CNO treatment, as exemplified in high magnification images. Scale bars: 100 µm (left), 10 µm (right). Plot of PNN density around PV+ cells ipsi- and contralateral to hM4Di expression normalized for each mouse to mean density in contralateral hemicortex. Indicated in bars are the number of cells analyzed in 4 CNO-treated and 3 PBS-treated mice. * Significantly different from other conditions.

Our results collectively suggest that PNN density is dynamically regulated in the adult depending on the activity level of PV interneurons.

### The PNN may be regulated cell-autonomously by each PV interneuron

During histochemical analyses of CNO treated mice, we noticed that hM4Di-negative PV interneurons had dense PNNs, whereas the PNN of neighboring hM4Di+ cells was reduced (see examples in Fig.5A). This suggests that each PV interneuron is able to regulate its own PNN cell-autonomously. In order to test this hypothesis, we compared PNN density within pairs of hM4Di+ and hM4Di-PV interneuron neighbors located at short distance from each other in the V1 area (maximal distance 106 µm, see Methods and Fig.5B). We verified that PNN density around hM4Di-cells was higher than around hM4Di+ cells (WFA staining intensity 85.1 ± 5.5 % of hM4Di-negative cells, n=21 cell pairs from 3 animals; Fig.5B). We next plotted WFA staining intensity of hM4Di-cells against that of hM4Di+ cells for each PV interneuron pair (Fig.5B). The plot was skewed towards higher WFA staining around hM4Di-cells, consistent with hM4Di effect on the PNN. Linear regression analysis yielded a slope of 0.2, which did not significantly differ from the zero slope value expected from independence between WFA staining intensities of hM4Di+/PV+ and hM4Di-/PV+ cells (p=0.12). These results suggest that PNN density is regulated by each PV interneuron independently of its PV cell neighbors.

**Figure 5.**
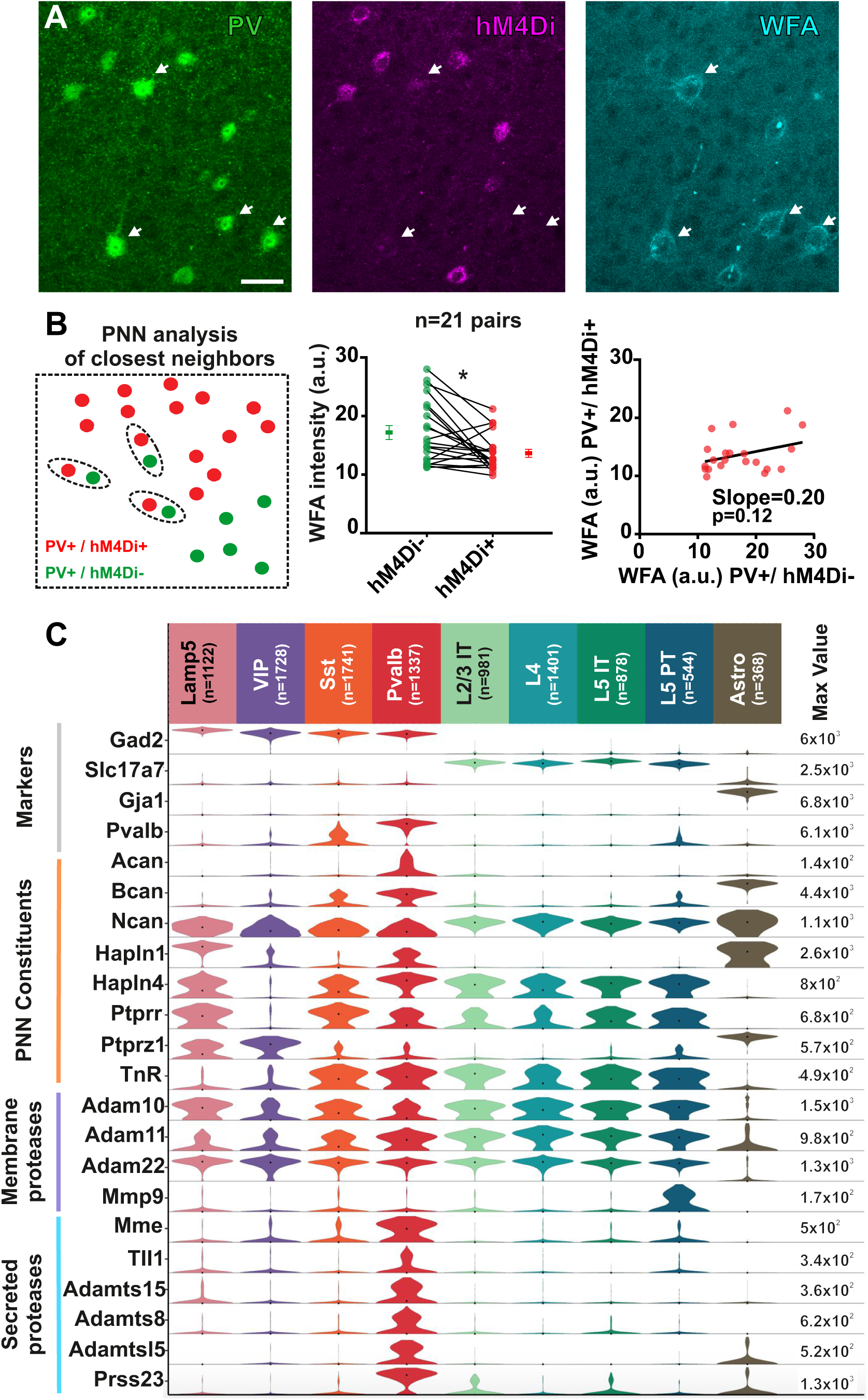
PV interneurons may regulate their PNN cell-autonomously. (**A**) Examples of the high PNN density (arrows) observed around hM4Di-negative, or weakly hM4Di-positive PV interneurons, as compared to the low PNN density observed around their PV+ neighbors robustly expressing hM4Di in the V1 cortex of a CNO treated mouse. Scale bar: 20 µm. (**B**) Comparison of PNN density within pairs of hM4Di+ and hM4Di-closest neighbors PV+ cells: selection criterion (left), individual and mean WFA intensity (middle, * significant difference) and scatter plot of WFA intensity for PV interneuron pairs (right, n=21). Note that the slope of linear regression did not significantly differ from zero. (**C**) Violin plots showing distributions of individual gene expression in 10,100 single cells from primary visual cortex of P56 mice. Data are from the Allen Institute. Cells are from subclasses segregated in Tasic et al. (20): GABAergic (Lamp5, Vip, Sst and Pvalb), glutamatergic (layers 2/3 and 5 IT intratelencephalic, layer 4, layer 5 PT Pyramidal Tract) and astrocytes. Rows show individual gene expression across cell types, values (number per million reads) are displayed on a log10 scale normalized to maximum expression value for each gene (right column), black dots are median values. *Markers*: Gad2, glutamate decarboxylase; Slc17a7, vesicular glutamate transporter; Gja1, gap junction alpha1; Pvalb, PV; *Lecticans*: Acan, aggrecan; Bcan, brevican; Ncan, neurocan; *PNN linkers*: Hapln, hyaluronan and proteoglycan link protein; Ptprr and Ptprz1, Protein Tyrosine Phosphatase Receptor types R and Z1; Tnr, tenascin R; *Proteases*: Adam, A Disintegrin And Metalloproteases; Mmp9, membrane metallopeptidase 9; Mme, neprilysin; Tll1, tolloid-like metallopeptidase; Adamts, A Disintegrin And Metalloprotease with Thrombospondin motif; Prss23, serine protease 23.

PV interneurons express multiple genes involved in synthesis and degradation of the PNN (Okaty et al., 2009; Rossier et al., 2015). We thus searched through published transcriptomic database of single cells collected from the primary visual cortex of P56 mice (Tasic et al., 2018) for the expression of genes that may enable PV interneurons to regulate their PNN cell-autonomously. Figure 5C shows expression levels of selected PNN lecticans, linkers, and proteases in 10,100 cells comprising major interneuron and layers II-V excitatory neuron types as previously defined (Tasic et al., 2018), as well as astrocytes. The expression profiles of PNN-related genes differed between cell-types, with the largest array of genes expressed by PV interneurons. All lecticans and PNN linkers mRNAs were present in PV interneurons, which preferentially express Acan and weakly express Ptprz1 mainly found in astrocytes (Rossier et al., 2015; Maurel et al., 1994). PV interneurons strikingly differed from other cell types in expressing significant levels of all tested proteases, except Mmp9 mainly observed in a subtype of layer V excitatory neurons. Notably, secreted proteases were preferentially expressed by PV interneurons consistent with earlier observations (Rossier et al., 2015). These results confirm that adult PV interneurons express key genes enabling cell-autonomous control of the accumulation and degradation of their own PNN.

## DISCUSSION

We tested the effects of targeted chemogenetic modulation of PV interneurons and excitatory neurons on PNN density around PV interneurons in the adult visual cortex. Inhibition of PV interneurons using the Gi-coupled hM4Di or the chloride channel PSAM-GlyR, as well as inhibition of glutamatergic neurons, induced PNN regression. Inhibition of PV interneurons did not elicit unbalanced excitation of the network and excitation of glutamatergic neurons or of PV interneurons did not alter PNN density, ruling out network disinhibition as a cause of PNN regression. We also found that the PNN of hM4Di-expressing PV cells was reduced compared to the PNN of their hM4Di-negative neighbors, and that PV interneurons express genes enabling control of their own PNN density. Our results indicate that silencing of PV interneurons, directly or through reduced synaptic excitation, triggers PNN regression in the adult cortex, and suggest that each PV cell regulates its own PNN cell-autonomously.

### Targeted chemogenetic modulation of PV interneurons or excitatory neurons

Cre-dependent viral transduction resulted in robust and selective expression of chemogenetic actuators, consistent with previous studies (Alexander et al., 2009; Hippenmeyer et al., 2005; Gorski et al., 2002; Magnus et al., 2011; Krashes et al., 2011). Cell-type-specific targeting was assessed by immunochemistry and patch-clamp recordings of transduced neurons, which all exhibited electrophysiological properties typical of PV interneurons or glutamatergic neurons (Cauli et al., 1997; Connors and Gutnick, 1990). Our results exclude DREADD-independent effects of CNO (Gomez et al., 2017) as the cause of PNN changes: (i) CNO was ineffective in hM3Dq expressing mice; (ii) CNO induced PNN regression in the hM4Di-expressing hemicortex but not contralaterally; (iii) CNO and PSEM treatments both effectively reduced PNN around PV interneurons expressing their cognate chemogenetic actuator; (iv) our electrophysiological recordings of acute CNO effects confirm its efficiency in modulating somatodendritic excitability of DREADD-expressing neurons (Alexander et al., 2009; Krashes et al., 2011 but see Stachniak et al., 2014; Gomez et al., 2017). Hence, the PNN changes we observed stem from specific chemogenetic ligand/actuator interactions eliciting a decrease in excitability of selectively targeted cell-types.

### PNN regression is triggered by electrical silencing of PV interneurons, but not by network disinhibition

Modulation of PV interneurons using hM4Di reinstates visual plasticity in the mouse cortex after closure of the critical period (Kuhlman et al., 2013). Consistent with our working hypothesis, we found that this chemogenetic paradigm also induced PNN regression. Our observations indicate that this effect, also obtained upon inhibition of PV cells by the chloride channel PSAM-GlyR, was due to electrical silencing of these interneurons. This outcome was reinforced by the reduced somatodendritic excitability of PV cells upon hM4Di activation, in agreement with earlier calcium imaging data (Kuhlman et al., 2009), and by the decrease in cortical gamma oscillations that critically rely on these interneurons (Cardin et al., 2009). Conversely, EEG recordings showed no evidence for unbalanced network excitation (e.g. epileptiform activities), and targeted excitation of either glutamatergic neurons or PV interneurons using hM3Dq did not alter PNN density, thereby ruling out network disinhibition as a cause of PNN regression. Our results further indicate that electrical silencing of PV interneurons triggers PNN regression in the adult independently of the activity of other network components. Indeed, both targeted inhibition of PV interneurons (which silences PV cells but disinhibits excitatory cells), and targeted inhibition of excitatory neurons (which silences both excitatory and PV cells), resulted in PNN regression. Interestingly, a reduction in PV interneuron function has been proposed to underlie the effects of several commonly prescribed drugs and of dark exposure on adult PNN density or cortical plasticity (Hensch and Quinlan, 2018).

Our results reveal that PNN remodeling in the adult relies on the ability of PV interneurons to sense their own activity level, which reflects the activity of their local and distant inputs. Indeed, PV interneurons receive strong inputs from sensory thalamic nuclei, are densely interconnected with excitatory neurons, and receive direct feedback on their own GABAergic activity through powerful autapses (Bacci et al., 2003; Gabernet et al., 2005; Faini et al., 2018). Among the candidate signals that may link neuronal activity of PV cells to PNN degradation, changes in post-synaptic calcium entry [e.g. through calcium permeable AMPA receptors (Geiger et al., 1995; Angulo et al., 1997)] are unlikely to be involved since PNN regression occurred regardless of excitatory neurons being inhibited or disinhibited. Conversely, a change in OTX2 import appears as a more likely candidate signal: OTX2 accumulation in PV interneurons is activity-dependent, and the reduction of OTX2 import induces PNN regression and reinstates juvenile plasticity in the adult (Sugiyama et al., 2008; Beurdeley et al., 2012).

### PV interneurons may regulate their PNN cell-autonomously

The present study provides evidence for cell-autonomous regulation of the PNN by each PV cell independently of its neighbors. This possibility is substantiated by analysis of transcriptomic data (Tasic et al., 2018) showing expression of a large array of genes that enable PV cells to control the density of their own PNN. The PNN regression observed in the present study can theoretically result from enhanced proteolysis or from reduced synthesis associated with tonic proteolysis. Reduced expression or secretion of lecticans or PNN linkers by individual PV cells is expected to reduce their PNN cell-autonomously. Moreover, the pattern of membrane and secreted proteases characteristically found in PV interneurons is indicative of a tonic level of PNN proteolysis, which may be upregulated at the level of gene expression, secretion or proteolytic activity upon PV interneuron silencing. Hence, the expression pattern of PNN-related genes observed in PV interneurons indicates that these cells are able to reduce their PNN cell-autonomously.

Multiple earlier studies indicate that PNN removal renders adult cortical network permissive for high circuit plasticity. The chemogenetic paradigm used in the present study is known to reinstate visual plasticity in the young adult after closure of the critical period (Kuhlman et al., 2013). Our results thus suggest that each PV cell is able to regulate the PNN-dependent plasticity of its microcircuit by local sensing of neuronal activities, thereby contributing to point-by-point tuning of cortical network properties in the adult.

## MATERIALS AND METHODS

### Animals, Viruses and Surgery

Experiments were carried out in accordance with the European Communities Council Directive 86/609/062, and animal protocols approved by our local ethics committee (Ce5/ 2012/062). Transgenic mice lines from Jackson laboratories: PV-Cre [# 008069, Pvalb^tm1(cre)Arbr^, (Hippenmeyer et al., 2005)], and Emx1-Cre [# 005628, Emx1^tm1(cre)Krj^, (Gorski et al., 2002)], were genotyped by PCR with following primers. PV-Cre: wild-type forward CAGAGCAGGCATGGTGACTA, wild-type reverse AGTACCAAGCAGGCAGGAGA, mutant forward, GCGGTCTGGCAGTAAAAACTATC, mutant reverse GTGAAACAGCATTGCTGTCACTT (wild-type: 500 bp, mutant: 100 bp); Emx1 Cre: wild-type forward AAGGTGTGGTTCCAGAATCG, wild-type reverse CTCTCCACCAGAAGGCTGAG, mutant forward GCGGTCTGGCAGTAAAAACTATC, mutant reverse GTGAAACAGCATTGCTGTCACTT (wild-type: 102 bp, mutant: 378 bp).

Adeno-associated pseudovirions (AAVs) encoding Designed Receptor Exclusively Activated by Designer Drug [DREADD, (Alexander et al., 2009)] AAV2/5-hSyn-DIO-hM4Di-mCherry (titer: 5.2×10^12^ gc/ml) and AAV2/5-hSyn-DIO-hM3Dq-mCherry (7.8×10^12^ gc/ml) were produced from Addgene plasmids #44362 and #44361 at the facility of Nantes University (UMR 1089, France). AAV2/5-hsyn-FLEX:rev-PSAM^L141F,Y115F^-GlyR-IRES-GFP (3.6×10^12^, diluted at 1×10^12^ gc/ml) was generously provided by Dr. C.J. Magnus [Sternson Lab, Janelia Research Campus, USA, (Magnus et al., 2011)].

For viral transduction, postnatal day (P) 25 to 28 PV-Cre or Emx1-Cre mice were anesthetized by intraperitoneal (i.p.) injection of ketamine/xylazine (100/10 mg/kg body weight) and restrained in a neonatal stereotaxic adaptor (David Kopf instrument). The scalp was retracted, and a burr hole was drilled in the skull at coordinates AP= 0.05 mm and ML=2 mm from lambda to target the V1 area of the right visual cortex. Viral suspension (0.5 µl for hM3Dq and hM4Di, 1 µl for PSAM-GlyR) was injected with a glass capillary (1 µm tip, Drummond) at 500 µm below the pial surface at a speed of 100 nl/min. The scalp was sutured and mice were housed for at least 4 weeks with food and water *ad libitum*.

### Chemogenetic treatment and histological processing

Four weeks after viral injection, DREADD-expressing mice received 4 i.p. injections at 12 h intervals of DREADD agonist Clozapine-N-oxide (CNO, 1mg/kg; HelloBio) or phosphate buffered saline (PBS: Na phosphate 10 mM, NaCl 137 mM, KCl 2.7 mM, pH 7.4; 100 µl) (Fig.S1). PSAM-GlyR-expressing mice were treated similarly with the PSAM agonist PSEM^89S^ (10 mg/kg, kind gift of Dr. C.J. Magnus, Magnus et al. 2011) or PBS.

One day after the last i.p. injection, mice were anesthetized using a lethal mix of ketamine/xylazine (200/20 mg/kg body weight, respectively) and perfused transcardially with PBS containing 4% paraformaldehyde. Brains were extracted, incubated 2 h at 4°C in the same fixative, and sectioned in 50 µm coronal slices using a vibratome (VT1000S, Leica). Free-floating sections were blocked for 1.5 h at room temperature in PBS/0.25% Triton X-100/ 0.2% gelatin solution (PBS-GT) and incubated overnight at 4°C in PBS-GT with primary antibodies against PV, and RFP (DREADD-mCherry) or GFP (PSAM-GlyR). Slices were then washed with PBS and incubated for 1.5 h at room temperature with relevant secondary antibodies in PBS-GT. After washing in PBS, slices were next incubated with biotinylated *Wisteria floribunda* Agglutinin (WFA, 10 mg/ml, CliniSciences) for PNN labeling. Slices were then washed and incubated with streptavidin-AMCA (1:1000, Vector Laboratories). Finally, slices were washed and mounted on gelatin-coated slides in Fluoromount-G (Southernbiotech). Antibodies were used at following dilutions: mice IgG1 anti-PV (1:1000, Sigma), rat anti-RFP (1:500, Chromotek), chicken anti-GFP (1:1000, Aves Labs), goat anti-mouse IgG1 Alexafluor488 (1:500, Life Technologies), goat anti-mouse IgG Alexafluor555 (1:500, Life Technologies), goat anti-rat IgG Alexafluor555 (1:500, Life Technologies), donkey anti-chicken IgY Alexafluor488 (1:400; Jackson Immunoresearch).

Fluorescence images were acquired using an epifluorescence macro-apotome (Axiozoomer, Zeiss) equipped with filters DAPI, GFP and CY3 to acquire images of entire sections, an epifluorescence microscope (DMR, Leica) equipped with filters A4, GFP and CY3 to analyze PNN density, and a laser scanning confocal microscope (SP5, Leica) with 20X, 40X and 63X objectives, and 405, 488, and 561 nm lasers. Images were processed using ImageJ (U.S. National Institutes of Health, Bethesda, MD, USA; http://rsbweb.nih.gov/ij/).

### PNN density analyses

PNN density was analyzed by quantifying WFA fluorescence intensity around PV immunoreactive cells in the V1 area of both ipsilateral (virally transduced) and contralateral (uninjected) visual cortices using wide-field microscopy. Only brains showing extended viral transduction in V1 were kept for further analysis. In the case of targeted transduction of PV interneurons, only mCherry+ (DREADD-expressing) or GFP+ (expressing PSAM-GlyR) cells were analyzed ipsilaterally. The region analyzed in V1 was selected using a 20X objective, based on conspicuously low WFA staining in V2 area [Fig.S1, (Ueno et al., 2018)], as delineated by Paxinos and Franklin (Paxinos and Franklin, 2004). Images were acquired in layers IV-V using a 63X objective, starting medially from the V2/V1 border and progressing laterally into V1 through contiguous acquisition fields. In each field, all PV+ (or ipsilateral PV+/mCherry+ or PV+/GFP+) cells were selected for PNN density analysis (Fig.S1). Images were acquired with constant brightness and contrast, and variable exposure time. Perisomatic PNN was delineated manually based on WFA staining intensity (Fig.S1), forming ring-shaped regions of interest (ROIs) defined using the XOR function of the FIJI software (ImageJ). For each ROI, we measured area and mean WFA fluorescence intensity, which was normalized for exposure time. For each animal, the average fluorescence intensity of contralateral ROIs was used to normalize fluorescence intensity of each ipsi- and contralateral ROI. For each animal, 3-4 bilateral sections were used for this analysis. Data obtained in different mice for a given condition were compared using a Kruskall-Wallis nonparametric test. Since no significant difference was observed between mice for a given condition, data were pooled. Between-group comparisons were performed using Mann–Whitney nonparametric test.

In order to investigate cell-autonomous regulation of the PNN, WFA staining intensity was compared within pairs of PV+ cells: one being hM4Di+ (mCherry+) and the other being its closest hM4Di-(mCherry-) neighbor. Confocal images were acquired with a 40X objective at the lateral edge of the hM4Di expressing zone within the V1 area, in order to maximize the number of hM4Di+/PV+ and hM4Di-/PV+ cell pairs. Pairs were selected based on a distance criterion: the mean of the minimal distance between hM4Di+/PV+ cells determined for each section [range: 70-106 µm, n=7 sections, N=3 mice] was used as the maximal radius to select pairs of hM4Di+ and hM4Di-PV+ neighboring cells. PNN density was quantified as described above, and compared within cell pairs. Comparison between hM4Di+/PV+ and hM4Di-/PV+ cells was performed using Wilcoxon nonparametric test. Linear regression analysis was performed to test the independence between WFA staining intensities of hM4Di+/PV+ and hM4Di-/PV+ cells.

### Electrophysiological recordings of CNO/DREADD effects in cortical slices

Four to ten weeks after viral injection, mice were anesthetized using a lethal mix of ketamine/xylazine (200/20 mg/kg body weight) and 250 µm-thick coronal slices of V1 visual cortex prepared for whole-cell patch-clamp recordings (Hay et al., 2018) performed on mCherry+ layers IV-V PV interneurons or layer V pyramidal cells selected under epifluorescence illumination with a 535-nm LED (CoolLED) and an RFP filter set (Semrock). Membrane potentials were not corrected for liquid junction potential. Cells were set at −60 mV by continuous current injection and submitted to series of current pulses (800 ms, from – 100 to 375 pA with 25 pA increments). Parameters were measured from three series of pulses in control conditions and 2 minutes after the beginning of CNO bath application. Resting potential was determined on sweeps where no current was injected, input resistance on responses to hyperpolarizing pulses eliciting 10-15 mV shifts, and rheobase defined as the minimal depolarizing pulse triggering an action potential. Between-group comparisons were performed using paired Wilcoxon nonparametric test.

### Electroencephalographic (EEG) recordings of CNO/DREADD effects *in vivo*

Four weeks after viral injection, mice were subjected to anesthesia induced with 2% isoflurane and maintained by ketamine/xylasine (100/10 mg per kg body weight), and their body temperature maintained at 36.5°C. Electrodes made of bundles of insulated tungsten wires were implanted through the skull and sealed in place with acrylic resin. An epidural screw placed above the olfactory bulb was used as ground. One electrode placed above the cerebellum was used as reference. Three recording electrodes were implanted ipsilateral to the hM4Di-expressing hemisphere in V1 (AP = 0.05 mm, ML = ± 2 mm, DV= −0.5 mm from lambda) and somatosensory S1 (AP = −2 mm, ML = 2 mm, DV = −0.5 mm from bregma) cortical areas. After a 1-week resting period, EEG and simultaneous video recordings of awake mice were performed as described (Sieu et al., 2015). A half hour recording was extracted in each condition for analysis using a custom-made software based on Labview (National Instruments). The EEG signal was filtered with 1-4 Hz, 6-10 Hz, 30-50 Hz, 55-95 Hz and 100-150 Hz pass-bands and mean power values were extracted for each band and normalized to control condition. Data obtained in different conditions were compared using a Student t-test.

Throughout this study, treatments (ligand vs. PBS) or experimental conditions (transduction of PV-Cre or Emx1-Cre with DREADDs- or PSAM-encoding AAVs) were allocated randomly across mice, and results are given as mean ± standard error of the mean (SEM), and a p-value below 0.05 was considered statistically significant.

## ACKNOWLEDGEMENTS

We thank Dr. Chris Magnus for the kind gift of chemogenetic PSAM-PSEM tools, the Allen Institute for sharing data and analysis tools, Drs. Maria Cecilia Angulo and Alberto Bacci for advice throughout this study, Bernadette Hanesse and the IBPS Cell Imaging and Animal Facilities for their valuable help.

---

**Fig. S1.**
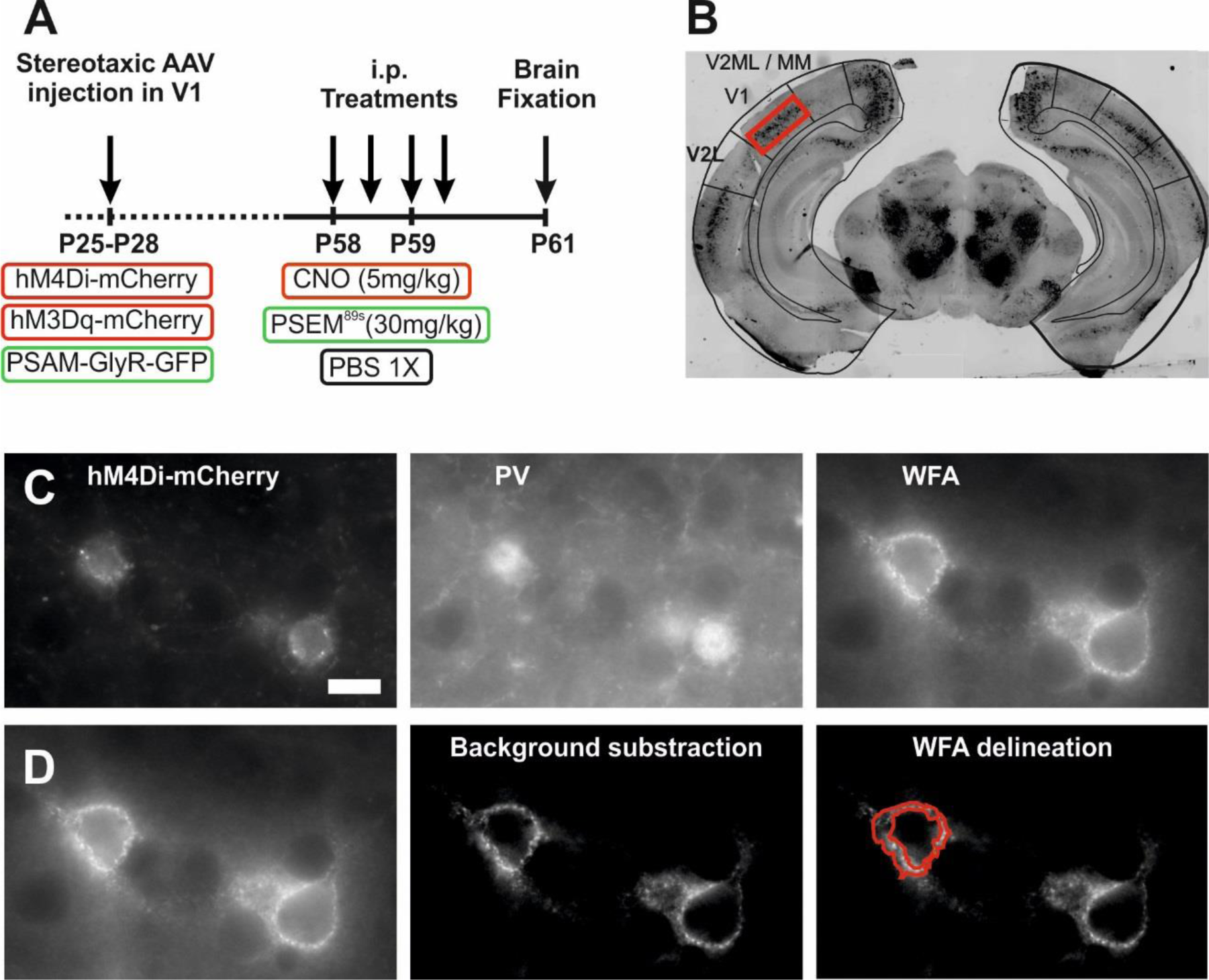
Chemogenetic paradigm and histological analysis of PNN density in the V1 visual cortex. (**A**) Four weeks after hemilateral AAV injection in the V1 cortex, DREADD- or PSAM-GlyR-expressing mice received 4 injections of relevant agonist or PBS starting at P58, and brains were processed for histochemistry at P61. (**B**) Macrotome fluorescence negative picture of a coronal section of mouse brain at the level of the visual cortex showing PNN staining with WFA. The superimposed section of the mouse brain atlas delineates the densely stained V1 area flanked by V2L and V2ML areas exhibiting faint PNN labeling. Throughout this study, analyses of PNN density were performed in layers IV-V of the V1 area as indicated by the red rectangle. (**C**) PNN density analyses were performed around PV immunopositive cells, or around cells showing expression of both PV and the chemogenetic tool, as exemplified here for the hM4Di-mCherry fusion protein. (**D**) In order to quantify PNN density, whole-field fluorescence pictures were acquired (left panel), background-subtracted using the Substract Background function of the ImageJ software (middle panel), and PNNs were delineated manually around the soma based on WFA staining intensity (red lines in right panel), to create ring-shaped ROIs using the XOR function of ImageJ. Scale bar for C-D: 20 µm.

**Fig. S2.**
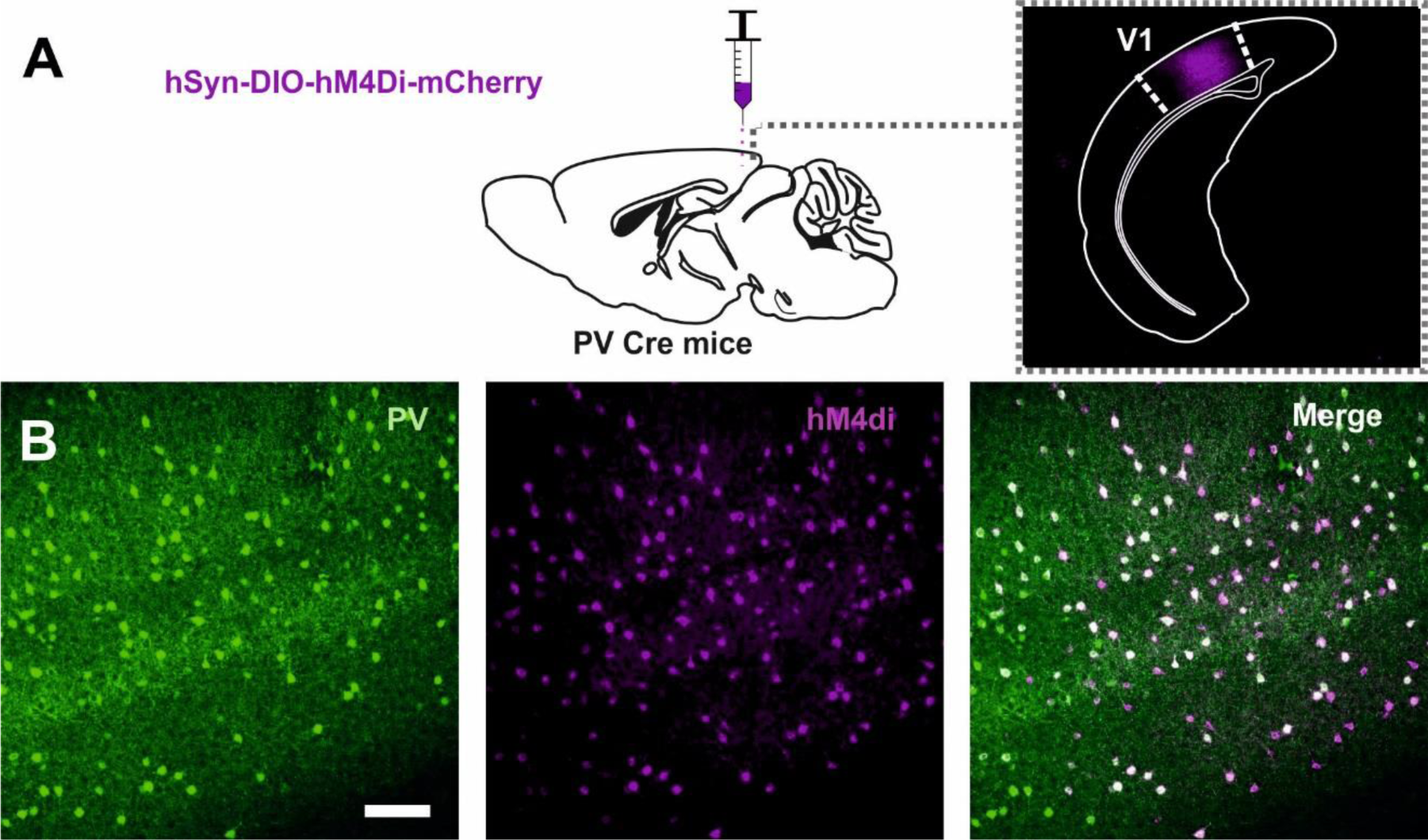
Targeted expression of DREADD hM4Di in PV interneurons. (**A**) Stereotaxic injection of Cre-dependent AAV encoding hM4Di fused to mCherry in the visual cortex of PV-Cre mice. Macrotome fluorescence picture showing expression of hM4Di-mCherry revealed by anti-RFP immunohistochemistry 5 weeks after injection. The superimposed section of the mouse brain atlas delineates the V1 area. (**B**) Confocal fluorescence images acquired in the V1 cortex after dual immunostaining showing hM4Di-mCherry expression in PV-positive cells. Scale bar: 100 µm.

**Fig. S3.**
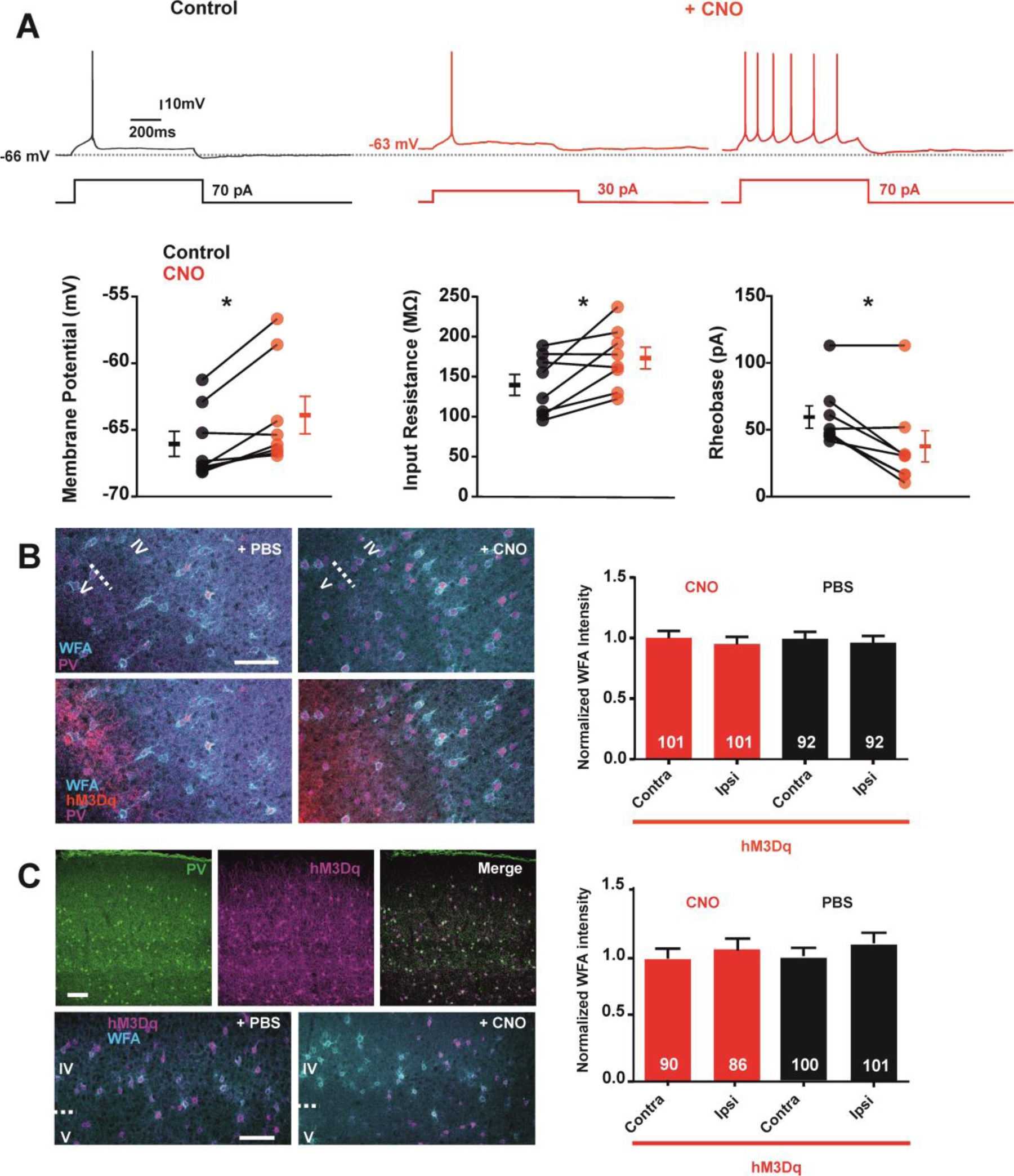
Targeted excitation of glutamatergic or PV neurons using hM3Dq does not alter PNN density. (**A**) Patch-clamp recordings in cortical slices. Traces show responses of a hM3Dq-expressing layer V pyramidal neuron to depolarizing current steps in control conditions and upon bath application of CNO (0.5 µM). CNO application elicited a depolarization of the membrane potential and a decrease of the current needed to induce action potential firing. Plots show electrophysiological parameters measured in control and CNO conditions in hM3Dq-expressing neurons (n=8). * Significant differences. (**B**) Confocal fluorescence images illustrate the PNN (WFA) surrounding PV+ cells in the vicinity of hM3Dq-expressing excitatory neurons in layers IV-V of the V1 cortex of mice treated with PBS or CNO. For better visualization, upper panels show WFA and anti-PV labelling separately from hM3Dq-mCherry positive excitatory neurons visible on lower panels. Scale bar: 100 µm. Plot of PNN density around PV+ cells ipsi- and contralateral to hM3Dq expression in the V1 cortex normalized for each mouse to mean density in contralateral hemicortex. Indicated in bars are the number of cells analyzed in 3 CNO-treated and 3 PBS-treated mice. (**C**) Upper confocal fluorescence images show hM3Dq-mCherry expression in PV+ cells of the V1 cortex 5 weeks after injection of corresponding Cre-dependent AAV in a PV-Cre mouse. Lower confocal fluorescence images illustrate the PNN (WFA) surrounding hM3Dq+ PV interneurons in layers IV-V after PBS or CNO treatment of the mice. Scale bars: 100 µm. Plot of PNN density around PV+ (contralateral uninjected hemicortex) and hM3Dq+/PV+ (ipsilateral injected) cells. Indicated in bars are the number of cells analyzed in 3 CNO-treated and 3 PBS-treated mice.

